# SCRIBE: a new approach to dropout imputation and batch effects correction for single-cell RNA-seq data

**DOI:** 10.1101/793463

**Authors:** Yiliang Zhang, Kexuan Liang, Molei Liu, Yue Li, Hao Ge, Hongyu Zhao

## Abstract

Single-cell RNA sequencing technologies are widely used in recent years as a powerful tool allowing the observation of gene expression at the resolution of single cells. Two of the major challenges in scRNA-seq data analysis are dropout events and batch effects. The inflation of zero(dropout rate) varies substantially across single cells. Evidence has shown that technical noise, including batch effects, explains a notable proportion of this cell-to-cell variation. To capture biological variation, it is necessary to quantify and remove technical variation. Here, we introduce SCRIBE (Single-Cell Recovery Imputation with Batch Effects), a principled framework that imputes dropout events and corrects batch effects simultaneously. We demonstrate, through real examples, that SCRIBE outperforms existing scRNA-seq data analysis tools in recovering cell-specific gene expression patterns, removing batch effects and retaining biological variation across cells. Our software is freely available online at https://github.com/YiliangTracyZhang/SCRIBE.

## 1 Introduction

Though undergoing rapid development in recent years, single-cell RNA-sequencing (scRNA-seq) data analysis is challenging due to severe systematic errors including batch effects and dropout events[1]. Batch effects may result from laboratory conditions, reagent lots and personnel differences, etc[2]. Combining data from multiple batches can offset the restriction of sample size and increase statistical power. However, the existence of batch effects makes data integration difficult since they are often associated with an outcome of interest[3]. Since the earliest observation of batch effects in microarray experiments[4], an explosion of batch effects correction methods for both bulk RNA-seq data[5–13] and scRNA-seq data[14–20] have been developed over the past two decades. Moreover, the inflation of zero caused by dropout events[21] renders more noise to scRNA-seq data than bulk RNA-seq data. Evidence has shown that batch effects account for a substantial percentage of dropout variability at the cell level[1]. Many imputation methods have been proposed in recent years to recover dropouts[18, 20, 22–25] in scRNA-seq data. However, among these methods, few take both batch effects and dropout events into consideration.

In this paper, we propose SCRIBE(Single-Cell Recovery Imputation with Batch Effects), a unified framework that jointly corrects batch effects and imputes dropout events across multiple biological groups. We apply SCRIBE to a single mouse neurons dataset to show that SCRIBE can accurately recover and correct batch effects for scRNA-seq data.

## 2 Method

### 2.1 SCRIBE model

We outline the framework of SCRIBE in this section. Let *Y*_*cg*_ denote the observed read count of gene *g* cell *c* and *Z*_*cg*_ be the dropout indicator. To account for dropout events and batch effects, we model *Y*_*cg*_ by a hierarchical zero-inflated Poisson mixed model[26, 27]:

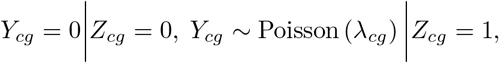

where 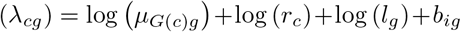. 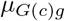 is the “true” normalized expression level for gene *g* of biological group *G*(*c*); *r*_*c*_ is the library size of cell *c*; *l_g_* is the total read count of gene *g* and *b*_*ig*_ is the random effect of batch *i* on gene *g*. We assume 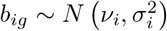 with a constraint ∑_*i*_ *ν*_*i*_ = 0 to ensure the identifiability of the parameters. *Z*_*cg*_ follows Bernoulli distribution with *P*(*Z*_*cg*_ = 1) = Φ [*γ*_*i*_ + *α*_*i*_ log (*r*_*c*_) + *β*_*i*_ log (*l*_*g*_)]. *γ*_*i*_, *α*_*i*_ and *β*_*i*_ are batch specific dropout parameters of batch *i* to which cell *c* belongs. Φ (·) is the CDF of standard normal distribution.

### 2.2 SCRIBE estimation procedure

The estimation of parameters in SCRIBE model follows maximum likelihood estimation(MLE). We apply Monte-Carlo EM(MCEM) algorithm[28, 29] here. Define *π*_*cg*_ = *γ*_*i*_ + *α*_*i*_ log (*r*_*c*_) + *β*_*i*_ log (*l*_*g*_). In order to apply data augmentation algorithm[30], we introduce a latent variable *η*_*cg*_ ~ *N*(*π*_*cg*_, 1) that satisfies 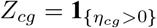. In MCEM, the random effect *b*_*ig*_ and the data augmentation variables *η*_*cg*_ are treated as missing data. In **E step**, with the help of Metropolis–Hastings algorithm[31, 32], we sample *b*_*ig*_ and *η*_*cg*_ for *K* times from the posterior distribution conditioning on the preceding **M step** estimation of other parameters. We denote the posterior samples as *b*_*igk*_, *η*_*cgk*_ and 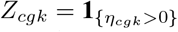, where *k* = {1, 2, …, *K*}. Then, in **M step**, the estimation of other parameters can be separated into three independent parts: (i) Estimation of 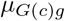. Directly using MLE estimator, we have 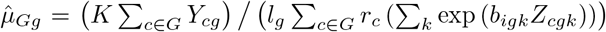; (ii) Estimation of *γ*_*i*_, *α*_*i*_ and *β*_*i*_. We fit a batch-specific linear model that regresses *η*_*cgk*_ on log (*r*_*c*_) and log (*l*_*g*_) to get the estimation 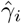, 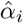 and 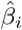; (iii) Estimation of *ν*_*i*_ and 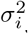 Restricted maximum likelihood(REML) is applied on *b*_*igk*_ to estimate 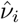 and 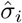 with the constrain ∑_*i*_*ν*_*i*_ = 0.

### 2.3 Dropout recovery imputation and batch effects correction

SCRIBE conducts data normalization, dropout imputation and batch effects correction simultaneously. Given the fitted model, define 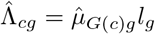 and let 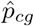 be the estimation of posterior probability, i.e. *p*_*cg*_ = *P*(*Z*_*cg*_ = 1|*Y*_*cg*_. To be specific, 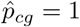, if *Y*_*cg*_ > 0. Otherwise, by Bayes’ theorem:

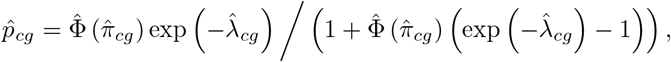

where 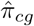 and 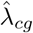 are the estimation of *π*_*cg*_ and *λ*_*cg*_ from the last iteration of MCEM, respectively. We develop a location and scale (L/S) adjustment[8] that retains the first and second moments of Poisson distribution. For *Y*_*cg*_ > 0, the corrected data 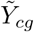 is given by

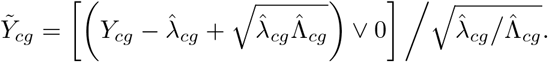

For *Y*_*cg*_ = 0, 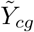 is the sum of corrected expression level and dropout imputation weighted by their posterior probability:

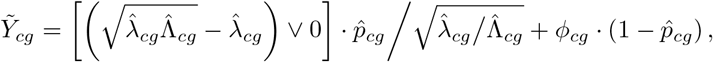

where 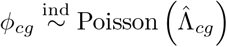.

## 3 Results

We use Usoskin’s single mouse neurons dataset[33] to examine the performance of SCRIBE on imputing the zeros, correcting batch effects and whether the imputation and batch effects correction would outperform existing analysis tools. The cell labels are assigned by the authors. We choose 4 cell types(peptidergic nociceptors, non-peptidergic nociceptors, neurofilament containing and tyrosine hydroxylase containing) with enough sample sizes from two batches. Genes with no expression are discarded. The filtered dataset contains 19,212 genes and 420 cells(265 cells in batch1 and 155 cells in batch2).

### 3.1 Comparison with existing dropout imputation methods

In this section, we benchmark SCRIBE with existing imputation methods including BUSseq[18], Drimpute[24], scImpute[22], SAVER[25] and MAGIC[23]. To generate realistic benchmarking datasets, we perform down-sampling experiments[25] on Usoskin dataset. We first select genes that have non-zero expression in at least 45% of the cells and cells with a library size of greater than 20,000 as the reference dataset, which we treat as the ‘true expression’ (data without dropouts). We end up with 3,156 genes and 244 cells(90 cells in batch1 and 154 cells in batch2) with overall detection rate 80.45%. Then, we randomly drop data in the reference dataset and generate an ‘observed dataset’. The probability of dropout is 0.8 for batch1 and 0.7 for batch2. In our simulation, we assume batch specific coefficients of library size *r_c_* and total read count *l_g_*, *α_i_* and *β_i_* are equal to zero for *i* = 1; 2. The overall detection rate is 21.15% after artificial dropout events which is close to 20.33%, the overall detection rate of Usoskin dataset.

All parameters are set to default for all the benchmarking imputation methods. To evaluate the performance of each method, we calculate the gene-wise Pearson correlation across cells and the cell-wise Pearson correlation across genes between the library-size normalized reference dataset and observed dataset, as well as between the normalized reference dataset and normalized observed dataset(Fig. 1a, b). The results show that SCRIBE outperforms other methods on both the gene-wise and cell-wise correlations although SAVER performs comparably to SCRIBE in terms of cell-wise correlations. Then, we calculate correlation matrix distance (CMD)[34] between gene-to-gene and cell-to-cell correlation matrices recovered by benchmarking methods as well as observed data and the reference gene-to-gene and cell-to-cell correlation matrix. CMD is a measure of the distance between two correlation matrices ranging from 0 (equal) to 1 (maximum difference). For gene-to-gene CMD(Fig. 1c), SCRIBE performs comparably to SAVER while it outperforms all other methods. For cell-to-cell CMD(Fig. 1d), SCRIBE significantly improves the performance.

**Figure 1:**
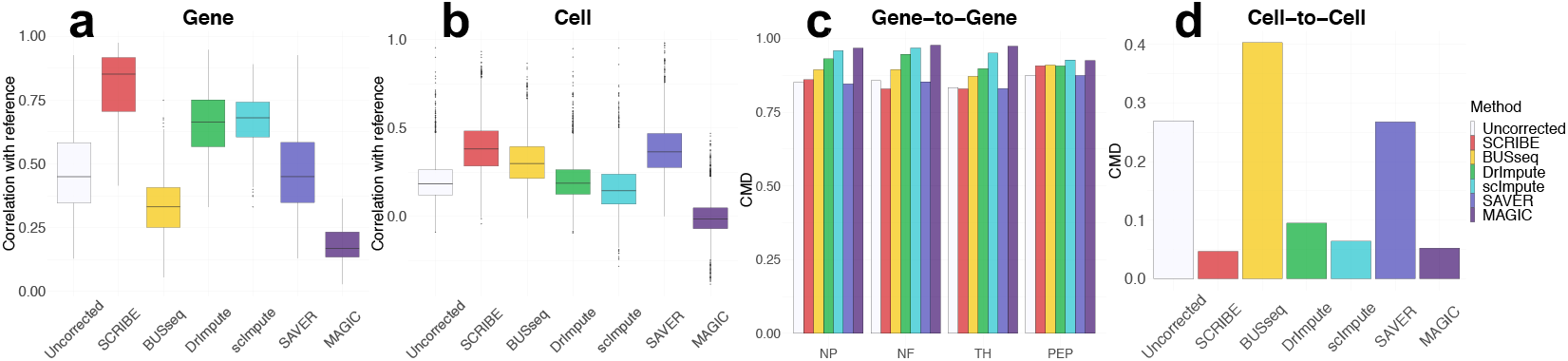
Comparisons of imputation methods on down-sampled single mouse neurons dataset. (**a**) Performance of methods measured by gene level correlation with reference. (**b**) Performance of methods measured by cell level correlation with reference. (**c**) Correlation matrix distance (CMD) between gene-to-gene correlation matrices recovered by benchmarking methods as well as observed data and the reference gene-to-gene correlation matrix. (**d**) Correlation matrix distance (CMD) between cell-to-cell correlation matrices recovered by benchmarking methods as well as observed data and the reference cell-to-cell correlation matrix.

### 3.2 Comparison with existing batch effects correction methods

Next, we benchmark SCRIBE with existing scRNA-seq batch effects correction methods including BUSseq[18], MNN[15], Seurat[14] and ZINB-WaVE[16]. The number of cell types is input to BUSseq with all other parameters set to default. The number of nearest neighbors *k* is set to 20 for MNN. All parameters in Seurat are set to default. The parameter of dimension *K* in ZINB-WaVE is set to 10. With the ground truth reported by the original publication, we use the Adjusted Rand Index (ARI) which measures the consistency between two clustering results as the performance metric. Higher ARI suggests better performance on batch effects correction. We implement k-means for cell types clustering on the results of each method except for BUSseq, which has its own inferred cell types. Our results of ARI are 0.930 for SCRIBE, 0.829 for BUSseq, 0.701 for MNN, 0.670 for Seurat and 0.904 for ZINB-WaVE. Uniform Manifold Approximation and Projection(UMAP)[35] plots show that SCRIBE outperforms all other methods in that SCRIBE can cluster the cells by cell types(Fig. 2). We also calculate the silhouette width[36] to compare the clustering performance of the different methods to remove batch effects and combine data. A high value of silhouette width indicates that the cell is well matched to its own cluster and poorly matched to neighboring clusters. Here, silhouette widths are calculated based on UMAP for either batch information or known sample group. Our results show that batch information after corrected by SCRIBE merges together and SCRIBE gives more concentration on high silhouette scores on cell types than other methods(Fig. 3).

**Figure 2:**
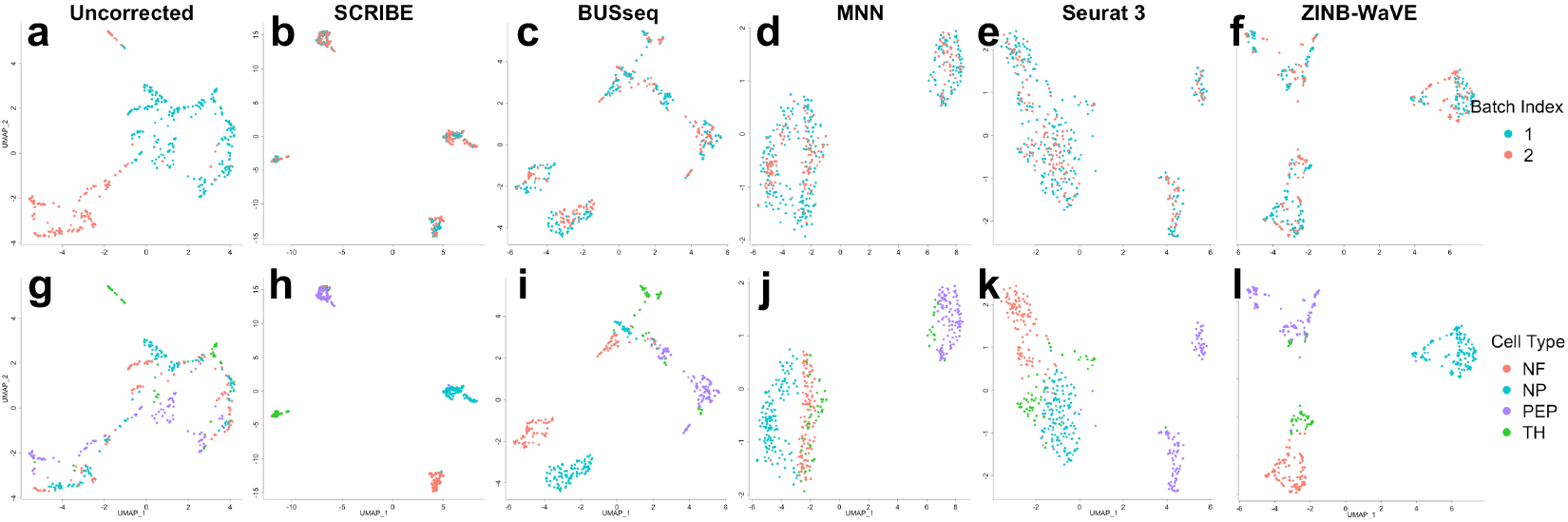
Comparisons of batch effects correction methods by Uniform Manifold Approximation and Projection(UMAP). **(a)** Uncorrected data colored by batch. **(b)** SCRIBE results colored by batch. **(c)** BUSseq results colored by batch. **(c)** MNN results colored by batch. **(d)** Seurat results colored by batch. **(e)** Seurat results colored by batch. **(f)** ZINB-WaVE results colored by batch. **(g)** Uncorrected data colored by cell types. **(h)** SCRIBE results colored by cell types. **(i)** BUSseq results colored by cell types. **(j)** MNN results colored by cell types. **(k)** Seurat results colored by cell types. **(l)** ZINB-WaVE results colored by cell types.

**Figure 3:**
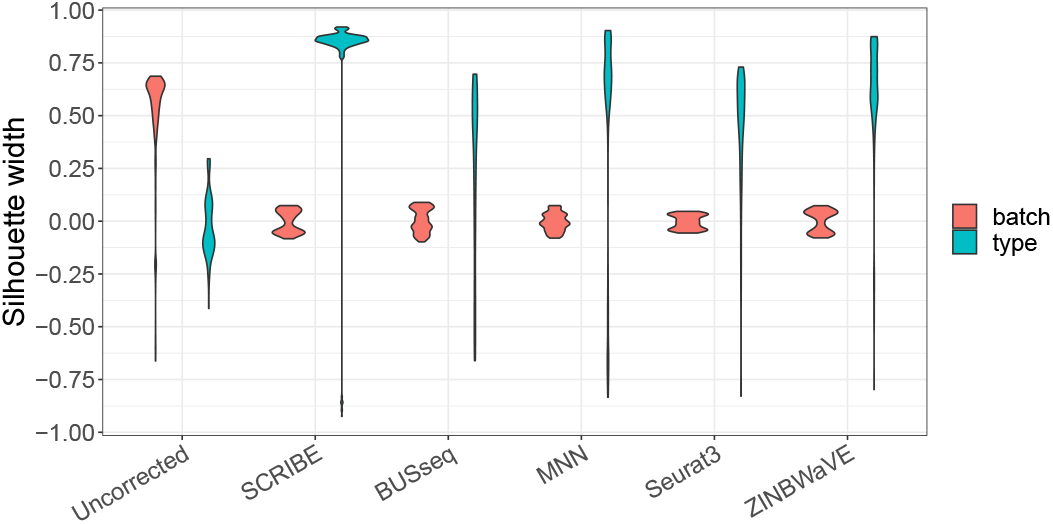
Silhouette width calculated on either batch information or known sample group for benchmarking methods on batch effects correction.

## 4 Conclusion

Although our understanding of scRNA-seq data is still far from complete, we have gained valuable knowledge from the technique. To make full use of this tool and capture biological insights behind the data, we propose a new statistical framework based on zero-inflated Poisson mixed model for scRNA-seq data analysis. Real data results show that SCRIBE is a powerful method that jointly adjusts for batch effects and imputes dropout events. For dropout imputation, SCRIBE borrows information from same cell types across other batches to recover the true expression level; For batch effects correction, random effect model in SCRIBE provides more robust adjustment for the batch effects on each gene as well as retaining biological variation.

### Future work

Directly following the current model, we can use the parameters and posterior distributions estimated by SCRIBE to develop a statistical inference procedure to test differential expression of markers across biological groups. New markers or cell types might be identified with this statistical testing procedure which will constitute the object of future studies.

## Acknowledgement

The authors would like to thank Dr. Yanyi Huang and Dr. Xiannian Zhang for their insightful suggestions.

## References

[1] Stephanie C Hicks, F William Townes, Mingxiang Teng, and Rafael A Irizarry. Missing data and technical variability in single-cell rna-sequencing experiments. Biostatistics, 19(4):562–578, 2017.

[2] Jeffrey T Leek, Robert B Scharpf, Héctor Corrada Bravo, David Simcha, Benjamin Langmead, W Evan Johnson, Donald Geman, Keith Baggerly, and Rafael A Irizarry. Tackling the widespread and critical impact of batch effects in high-throughput data. Nature Reviews Genetics, 11(10):733, 2010.

[3] Joshua M Akey, Shameek Biswas, Jeffrey T Leek, and John D Storey. On the design and analysis of gene expression studies in human populations. Nature genetics, 39(7):807, 2007.

[4] Eric S Lander. Array of hope. Nature genetics, 21(1s):3, 1999.

[5] Orly Alter, Patrick O. Brown, and David Botstein. Singular value decompositin for genome-wide expression data processing and modeling. Proceedings of the National Academy of Sciences of the United States of America, 97(18):10101–6, 2000.

[6] Torsten O Nielsen, Rob B West, Sabine C Linn, Orly Alter, Margaret A Knowling, John X O’Connell, Shirley Zhu, Mike Fero, Gavin Sherlock, and Jonathan R Pollack. Molecular characterisation of soft tissue tumours: a gene expression study. Lancet, 359(9314):1301–7, 2002.

[7] Monica Benito, Joel Parker, Quan Du, Junyuan Wu, Xiang Dong, Charles M. Perou, and J. S. Marron. Adjustment of systematic microarray data bases. Bioinformatics, 20(1):105–114, 2004.

[8] W Evan Johnson, Cheng Li, and Ariel Rabinovic. Adjusting batch effects in microarray expression data using empirical bayes methods. Biostatistics, 8(1):118–127, 2007.

[9] Sarah E. Reese, Kellie J. Archer, Terry M. Therneau, Elizabeth J. Atkinson, Celine M. Vachon, Mariza De Andrade, Jean Pierre A. Kocher, and Jeanette E. Eckelpassow. A new statistic for identifying batch effects in high-throughput genomic data that uses guided principal component analysis. Bioinformatics, 29(22):2877–83, 2011.

[10] Jeffrey T Leek and John D Storey. Capturing heterogeneity in gene expression studies by surrogate variable analysis. PLoS Genet, 3(9):e161, 2007.

[11] Jung Ae Lee, Kevin K. Dobbin, and Jeongyoun Ahn. Covariance adjustment for batch effect in gene expression data. Statistics in Medicine, 33(15):2681–95, 2014.

[12] Andrey A Shabalin, Håkon Tjelmeland, Cheng Fan, Charles M Perou, and Andrew B Nobel. Merging two gene-expression studies via cross-platform normalization. Bioinformatics, 24(9): 1154–1160, 2008.

[13] Xiangyu Luo and Yingying Wei. Batch effects correction with unknown subtypes. Journal of the American Statistical Association, 114(526):581–594, 2019.

[14] Andrew Butler, Paul Hoffman, Peter Smibert, Efthymia Papalexi, and Rahul Satija. Integrating single-cell transcriptomic data across different conditions, technologies, and species. Nature biotechnology, 36(5):411, 2018.

[15] Laleh Haghverdi, Aaron TL Lun, Michael D Morgan, and John C Marioni. Batch effects in single-cell rna-sequencing data are corrected by matching mutual nearest neighbors. Nature biotechnology, 36(5):421, 2018.

[16] Davide Risso, Fanny Perraudeau, Svetlana Gribkova, Sandrine Dudoit, and Jean-Philippe Vert. A general and flexible method for signal extraction from single-cell rna-seq data. Nature communications, 9(1):284, 2018.

[17] Emma Pierson and Christopher Yau. Zifa: Dimensionality reduction for zero-inflated single-cell gene expression analysis. Genome biology, 16(1):241, 2015.

[18] Fangda Song, Ga Ming Chan, and Yingying Wei. Flexible experimental designs for valid single-cell rna-sequencing experiments allowing batch effects correction. bioRxiv, page 533372, 2019.

[19] Cheng Jia, Yu Hu, Derek Kelly, Junhyong Kim, Mingyao Li, and Nancy R Zhang. Accounting for technical noise in differential expression analysis of single-cell rna sequencing data. Nucleic acids research, 45(19):10978–10988, 2017.

[20] Romain Lopez, Jeffrey Regier, Michael B Cole, Michael I Jordan, and Nir Yosef. Deep generative modeling for single-cell transcriptomics. Nature methods, 15(12):1053, 2018.

[21] Peter V Kharchenko, Lev Silberstein, and David T Scadden. Bayesian approach to single-cell differential expression analysis. Nature methods, 11(7):740, 2014.

[22] Wei Vivian Li and Jingyi Jessica Li. An accurate and robust imputation method scimpute for single-cell rna-seq data. Nature communications, 9(1):997, 2018.

[23] David van Dijk, Juozas Nainys, Roshan Sharma, Pooja Kathail, Ambrose J Carr, Kevin R Moon, Linas Mazutis, Guy Wolf, Smita Krishnaswamy, and Dana Pe’er. Magic: A diffusion-based imputation method reveals gene-gene interactions in single-cell rna-sequencing data. BioRxiv, page 111591, 2017.

[24] Wuming Gong, Il-Youp Kwak, Pruthvi Pota, Naoko Koyano-Nakagawa, and Daniel J Garry. Drimpute: imputing dropout events in single cell rna sequencing data. BMC bioinformatics, 19(1):220, 2018.

[25] Mo Huang, Jingshu Wang, Eduardo Torre, Hannah Dueck, Sydney Shaffer, Roberto Bonasio, John I Murray, Arjun Raj, Mingyao Li, and Nancy R Zhang. Saver: gene expression recovery for single-cell rna sequencing. Nature methods, 15(7):539, 2018.

[26] Diane Lambert. Zero-inflated poisson regression, with an application to defects in manufacturing. Technometrics, 34(1):1–14, 1992.

[27] Daniel B Hall. Zero-inflated poisson and binomial regression with random effects: a case study. Biometrics, 56(4):1030–1039, 2000.

[28] James G Booth and James P Hobert. Maximizing generalized linear mixed model likelihoods with an automated monte carlo em algorithm. Journal of the Royal Statistical Society: Series B (Statistical Methodology), 61(1):265–285, 1999.

[29] Greg CG Wei and Martin A Tanner. A monte carlo implementation of the em algorithm and the poor man’s data augmentation algorithms. Journal of the American statistical Association, 85(411):699–704, 1990.

[30] Martin A Tanner and Wing Hung Wong. The calculation of posterior distributions by data augmentation. Journal of the American statistical Association, 82(398):528–540, 1987.

[31] Nicholas Metropolis, Arianna W Rosenbluth, Marshall N Rosenbluth, Augusta H Teller, and Edward Teller. Equation of state calculations by fast computing machines. The journal of chemical physics, 21(6):1087–1092, 1953.

[32] W Keith Hastings. Monte carlo sampling methods using markov chains and their applications. 1970.

[33] Dmitry Usoskin, Alessandro Furlan, Saiful Islam, Hind Abdo, Peter Lönnerberg, Daohua Lou, Jens Hjerling-Leffler, Jesper Haeggström, Olga Kharchenko, Peter V Kharchenko, et al. Unbiased classification of sensory neuron types by large-scale single-cell rna sequencing. Nature neuroscience, 18(1):145, 2015.

[34] Markus Herdin, Nicolai Czink, Hüseyin Ozcelik, and Ernst Bonek. Correlation matrix distance, a meaningful measure for evaluation of non-stationary mimo channels. In 2005 IEEE 61st Vehicular Technology Conference, volume 1, pages 136–140. IEEE, 2005.

[35] Leland McInnes, John Healy, and James Melville. Umap: Uniform manifold approximation and projection for dimension reduction. arXiv preprint arXiv:1802.03426, 2018.

[36] Peter J Rousseeuw. Silhouettes: a graphical aid to the interpretation and validation of cluster analysis. Journal of computational and applied mathematics, 20:53–65, 1987.

